# Crosstalk between repair pathways elicits Double-Strand Breaks in alkylated DNA: implications for the action of temozolomide

**DOI:** 10.1101/2020.11.21.391326

**Authors:** Robert P. Fuchs, Asako Isogawa, Joao A. Paulo, Kazumitsu Onizuka, Tatsuro Takahashi, Ravindra Amunugama, Julien Duxin, Shingo Fujii

## Abstract

Temozolomide (TMZ), a DNA methylating agent, is the primary chemotherapeutic drug used in glioblastoma treatment. TMZ induces mostly N-alkylation adducts (N7-methylguanine and N3-methyladenine) and some O^6^-methylguanine (O^6^mG). Current models propose that during DNA replication, thymine is incorporated across from O^6^mG, promoting a futile cycle of mismatch repair (MMR) that leads to DNA double strand breaks (DSBs). To revisit the mechanism of O^6^mG processing, we reacted plasmid DNA with N-Methyl-N-nitrosourea (MNU), a temozolomide mimic, and incubated it in Xenopus egg extracts. We show that in this system, mismatch repair (MMR) proteins are enriched on MNU-treated DNA and we observe robust, MMR-dependent, repair synthesis. Our evidence also suggests that MMR, initiated at O^6^mG:C sites, is strongly stimulated in *cis* by repair processing of other lesions, such as N-alkylation adducts. Importantly, MNU-treated plasmids display DSBs in extracts, the frequency of which increased linearly with the square of alkylation dose. We suggest that DSBs result from two independent repair processes, one involving MMR at O^6^mG:C sites and the other involving BER acting at a nearby N-alkylation adducts. We propose a new, replication-independent mechanism of action of TMZ, that operates in addition to the well-studied cell cycle dependent mode of action.

## Introduction

Alkylating agents, a class of important environmental carcinogens, have been widely used in molecular biology to study fundamental repair processes and in the clinic to treat cancer patients. Among the DNA adducts produced by methylating agents such as N-methyl-N-nitrosourea (MNU) and temozolomide (TMZ), a clinically used mimic, the most abundant are two N-alkylation adducts, at the N7 position of guanine (7mG: 70-75% of total alkyl adducts) and the N3 position of adenine (3mA: 8-12%). Importantly, both reagents also produce 8-9% O-alkylation adducts in the form of O^6^-methylguanine (O^6^mG). This feature contrasts with another common methylating agent, methyl-methane sulfonate (MMS), which forms a much lower level of O^6^mG (<0.3%) while producing similarly high proportions of 7mG (81-83%) and 3mA (10-11%)(Beranek, 1990). For many years, the differences in O versus N reactivities have been rationalized by differences in chemical reaction mechanisms; on one side, compounds such as MMS, with very low O-reactivity were classified as an SN2 agent (bimolecular nucleophilic substitution) while other agents, such as MNU and TMZ, with increased O adduct formation were called SN1 agents (monomolecular nucleophilic substitution). While this classification turned out not to be mechanistically accurate (Loechler, 1994), we will nevertheless use this nomenclature throughout this paper for the sake of simplicity. The major N-alkylation (N-alkyl) adducts (7mG and 3mA) are repaired by base excision repair (BER), using N-methylpurine-DNA glycosylase (MPG) also known as 3-alkyladenine DNA glycosylase (AAG) and alkylpurine DNA N-glycosylase (APNG) (Chakravarti et al., 1991; Lindahl, 1976). O-alkylation adducts (O^6^mG, O^4^mT) can be directly repaired by O^6^-methylguanine-DNA methyl transferase (MGMT), a protein that transfers the methyl group from these adducts to one of its cysteine residues (Demple et al., 1982; Olsson and Lindahl, 1980; Tano et al., 1990). In addition, alkylating agents also produce a variety of other minor (1-2%) N-alkyl adducts, namely 1mA, 3mC, 3mT and 1mG that are directly demethylated by AlkB homologs (Aas et al., 2003; Duncan et al., 2002; Falnes et al., 2002; Trewick et al., 2002). In summary, SN1 and SN2 alkylating agents produce a diverse array of DNA adducts, but they differ greatly in the amount of O^6^mG produced.

Agents such as MMS mostly induce N-alkyl adducts that lead to DSBs during S-phase as a consequence of BER repair. Indeed, inactivation of the AAG glycosylase, the BER initiating enzyme, suppresses DSB while inactivation of Pol β leads to their exacerbation (Simonelli et al., 2017; Tang et al., 2011; Trivedi et al., 2005). In rodent cells, it was proposed that MMS-induced DSBs arise when replication meets BER-induced single-strand breaks (Ensminger et al., 2014). The toxicity of N-alkyl adducts was found to depend on the cell type. AAG-mediated repair of N-alkyl adducts was found to mitigate toxicity in mouse ES cells and HeLa cells, while repair was shown to cause toxic intermediates in retina and bone marrow cells (Meira et al., 2009). In all cell type, *O*-alkyl adducts were found to be highly cytotoxic and mutagenic. While the mutagenicity of O^6^mG is easily accounted for by its high propensity to mispair with T during DNA synthesis (Bhanot and Ray, 1986; Loechler et al., 1984; Mazon et al., 2010) its cytotoxicity is intriguing since O^6^mG *per se* does not interfere with DNA synthesis. A seminal paper, published 50 years ago by Plant and Roberts (Plant and Roberts, 1971), noted that when synchronized HeLa cells are treated in G1 with MNU, they continue through the first cell cycle almost normally and with little effect on DNA synthesis. On the other hand, there is a dramatic effect on DNA synthesis in the second cell cycle after MNU exposure. These data led the authors to surmise that cytotoxicity stems from a secondary lesion that forms when DNA synthesis occurs across O^6^mG template adducts (Plant and Roberts, 1971). It was demonstrated later that MNU-mediated inhibition of DNA synthesis, in the first and second cycle, is due to the action of the mismatch repair machinery (MMR) that acts on O^6^mG:T lesions that form upon DNA synthesis (Noonan et al., 2012; Plant and Roberts, 1971; Quiros et al., 2010). (Goldmacher et al., 1986; Kat et al., 1993). Indeed, O^6^mG:T lesions were found to be excellent substrates for MMR (Duckett et al., 1999; Yoshioka et al., 2006). During MMR gap-filling, the O^6^mG:T mispair is reformed, potentially leading to another round of MMR, thus entering so-called futile MMR cycles (Kaina et al., 2007; Karran and Bignami, 1994; Olivera Harris et al., 2015; York and Modrich, 2006). The MMR cycling model has received experimental support *in vitro* (York and Modrich, 2006) and in *E. coli* (Mazon et al., 2010). The cytotoxic mode of action of alkylating agents that form O^6^mG adducts is believed to result from the encounter, by the replication fork, of these MMR intermediates leading to subsequent double-strand breaks (DSB), apoptosis and cell death (Ochs and Kaina, 2000). Studies with synchronized cells have shown that the critical events related to cytotoxicity occur in the 2nd cell cycle post treatment (Quiros et al., 2010). However, as discussed in recent review articles, the precise mechanism by which MMR leads to DSBs has yet to be established (Gupta and Heinen, 2019; Kaina and Christmann, 2019).

While most studies have been devoted to MNU-induced cell cycle effects, in the present paper we wanted to investigate the early response to MNU treatment, i.e. in the absence of replication. We addressed this question using Xenopus egg extracts, which recapitulate most forms of DNA repair (Wühr et al., 2014). Upon incubation in these extracts, plasmids treated with MNU, exhibit robust repair synthesis in the absence of replication. Repair synthesis occurs at O^6^mG:C lesions, depends on MMR, and involves an excision tract of several hundred nucleotides. MMR events at O^6^mG:C sites are robustly stimulated by additional processing at N-alkylation lesions, most likely via BER. Previous studies have described activation of MMR in the absence of replication in cells treated by SN1-methylating agents, a process termed noncanonical MMR (ncMMR) (Peña-Diaz et al., 2012). Interestingly, we observed replication-independent induction of DSBs in MNU-treated plasmids. The kinetics of DSB formation obeys a quadratic MNU dose-response suggesting the involvement of two independent repair events. We propose that DSBs occur when the gap generated at a O^6^mG adduct during MMR overlaps with a BER intermediate initiated at a N-alkyl adducts in the opposite strand. These data reveal a novel facet of MNU-induced damage to DNA that is replication independent. Extrapolation of the *in vitro* data led us estimate that ≈ 10 DSBs per cell can be induced by a single daily dose of TMZ used in the clinic in the absence of replication.

## Results

### 1. Identification of the proteins that specifically bind to DNA alkylation damage in Xenopus egg extracts

In order to identify proteins binding to O^6^mG-containing base pairs in Xenopus egg extracts, we used a recently developed plasmid pull-down procedure (Isogawa et al., 2020; 2018). To this end, plasmids were treated with MMS, which forms very low levels of O^6^mG, and with MNU or N-ethyl-N-nitrosourea (ENU). As SN1 agents, the latter two generate much more O-alk adducts (Beranek, 1990). These agents react chemically with DNA under neutral pH conditions, and we established *in vitro* reaction conditions that trigger comparable levels of plasmid alkylation (Fig S1A). The pull-down procedure involves immobilization of plasmid DNA on magnetic beads by means of a triple-helix forming probe (Fig. 1A) (Isogawa et al., 2020; 2018). The same amount of untreated or alkylated plasmids was coupled to magnetic beads and incubated in nucleoplasmic extracts (NPE) derived from Xenopus eggs (Walter et al., 1998). The reaction was stopped by dilution into a formaldehyde-containing buffer, which fixes protein-DNA complexes. After washing the beads and reversing the cross-links, the recovered proteins were visualized by silver staining following SDS-PAGE. As a negative control, beads with no DNA recovered only a low amount of protein illustrating efficient removal of unspecific proteins (Fig. S1B). Proteins captured on the different plasmids were analyzed by label-free mass spectrometry as described in Material and Methods. As shown in Fig. 1B, MMR proteins (labeled in red) were highly enriched on MNU-treated plasmid compared to undamaged control plasmid (Fig. 1B). All six canonical MMR proteins (MSH2, MSH3, MSH6, MLH1, PMS1 and PMS2) that comprise the MutSα, MutSβ, MutLα, and MutLβ heterodimers were recovered (Jiricny, 2006). RAD18, POLη, EXO1 and two subunits of Pol delta (POLD2 and POLD3) were also specifically enriched on MNU-plasmid. It was previously noted that upon oxidative stress, produced by hydrogen peroxide treatment, RAD18 and Polη are recruited to chromatin in a MSH2-MSH6 (MutSα) dependent manner in human cells (Zlatanou et al., 2011). While MutSα, MutSβ, and MutLα functionaly participate in MMR, the role of MutLβ (MLH1-PMS1) remains unknown (Jiricny, 2006). No MMR proteins were enriched on MMS-treated plasmids (Fig. 1C). As MNU treatment induces 20-30 times more O^6^mG adducts than MMS, we postulate that recruitment of MMR proteins depends on O^6^mG. Comparison of proteins captured on MNU-versus MMS-treated plasmids indeed reveals specific enrichment of MMR proteins. There is a concomitant loss of proteins specifically recruited at N-alkyl adducts since these adducts are equally present in MMS and MNU treated plasmids (Fig. S1C).

**Fig. 1.**
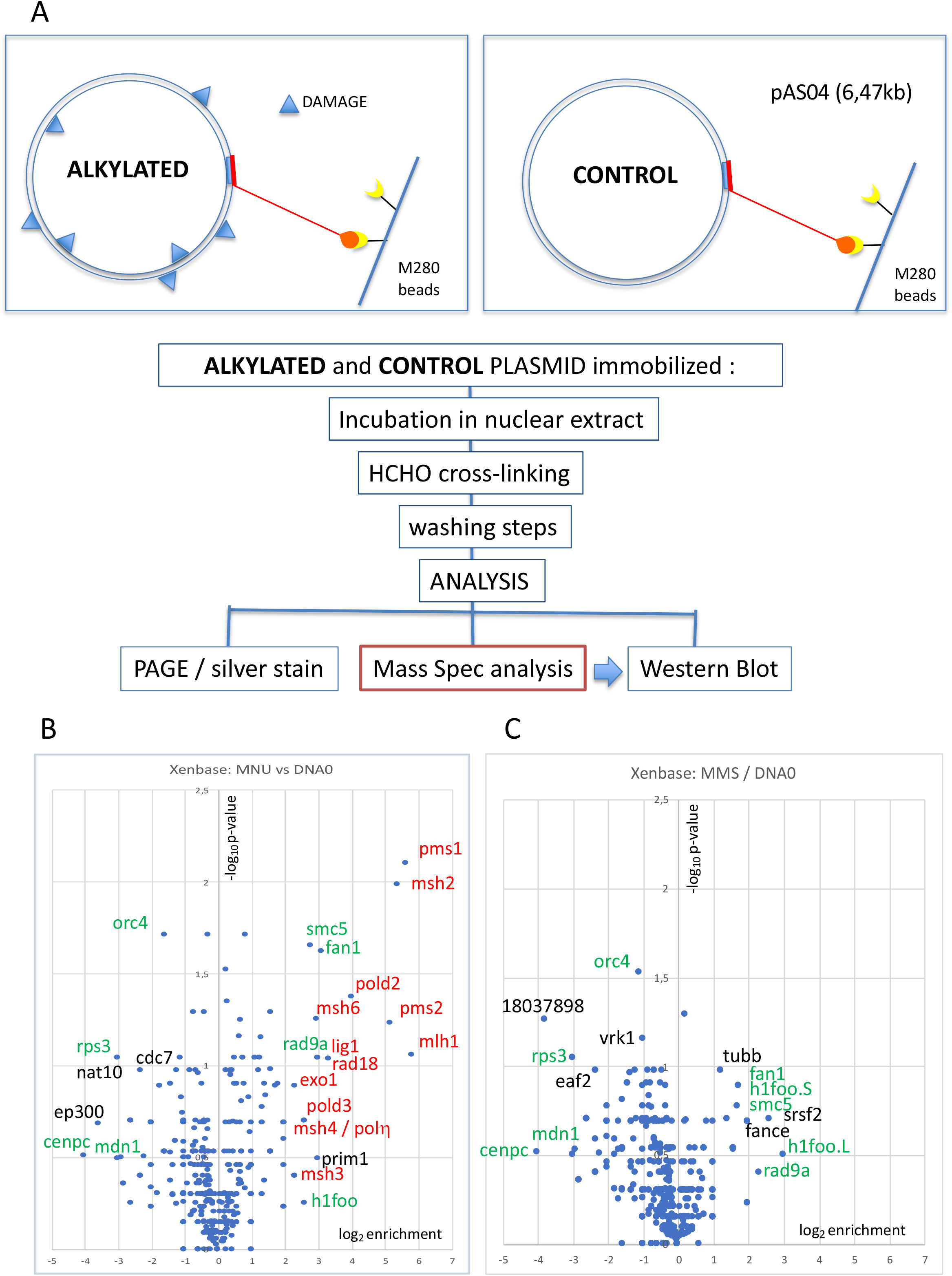
Pull-down of proteins that bind to alkylated-versus untreated-plasmid DNA. **A**. Experimental workflow. plasmid DNA (pAS04, 6.5kb) was treated with MMS or MNU under conditions leading to a similar extent of N-alkylation (≈ one alkaline cleavage site every 500 nt, see in Fig. 1SA). Immobilized plasmid DNA were incubated in Xenopus egg extract (NPE) for 10 min at room temperature under mild agitation. The reaction was stopped by addition of formaldehyde (0.8% final) to cross-link the proteins-DNA complexes. The beads were processed and analyzed by PAGE or by MS as described in Material and Methods. **B**. Relative abundance of proteins captured on MNU-treated versus untreated DNA0. Proteins captured on equal amounts of MNU- or untreated plasmid were analyzed by label-free MS in triplicates. For all proteins, average spectral count values in the MNU-treated plasmid sample were divided by the average spectral count values in the DNA0 sample. The resulting ratio is plotted as its log2 value along x-axis. The statistical significance of the data is estimated by the p value in the Student test and plotted as the -log10p along y-axis. Proteins enriched on MNU versus untreated plasmid DNA appear in the right-side top corner and essentially turn out to be MMR proteins labelled in red (fig 1B). Data shown are analyzed using Xenbase data base. **C**. Relative abundance of proteins captured on MMS-treated versus untreated DNA0. Proteins captured on equal amounts of MMS- or untreated plasmid were analyzed by label-free MS in triplicates. The data are analyzed and plotted as in panel B for MNU using Xenbase data base. Proteins (labeled in green in Fig. 1B and 1C) are found enriched or excluded in both MMS versus DNA0 and MNU versus DNA0 plasmids. We suggest these proteins are recruited or excluded from binding to DNA by the abundant class of N-alkylation adducts that both MMS- and MNU-treated plasmids share in common (∼27 N-alkyl adducts per plasmid).

In addition, compared to lesion-free control plasmid, some proteins were enriched on or excluded from both MMS and MNU-treated plasmids (Fig. 1B and 1C, green labels). We suggest that the recruitment or exclusion of these proteins depends on the abundant 7mG and 3mA adducts formed by both MMS and MNU. The reason why BER proteins, normally involved in the repair of these N-alkyl adducts, were not captured is unclear. One possibility is that BER proteins interact too transiently with DNA to be efficiently captured.

### 2. Repair of alkylated plasmid DNA in Xenopus egg extracts

We next investigated the repair of DNA treated by the different alkylating agents in Xenopus egg extracts. Plasmid was alkylated with MMS, MNU, or ENU to a density of on average one lesion every ≈500 nt (Fig. S1A). The alkylated plasmids were incubated in NPE in the presence of α^32^P-dATP. These extracts contain high levels of geminin, an inhibitor of replication licensing. Therefore, any observed DNA synthesis occurs independently of DNA replication and corresponds to so-called “unscheduled DNA Synthesis” (UDS)(Fig. 2A). Undamaged plasmid exhibited a low level of background DNA synthesis, whereas MNU and ENU treated plasmids sustained robust, time dependent UDS equivalent to 3-4% of the synthesis needed for a full round of replication (Fig. 2B). MMS-treated plasmid exhibited UDS that was just two-fold above the background seen in undamaged plasmid (Fig. 2B). Given that the assay measures incorporation of α^32^P-dATP, long patch BER events (Sattler et al., 2003) will be detected, while short patch BER events (1 nt patch) will only be detected at 3mA but not at 7mG adducts. The assay is clearly biased towards the detection of events such as MMR that involve repair patches hundreds of nucleotides long.

**Fig. 2.**
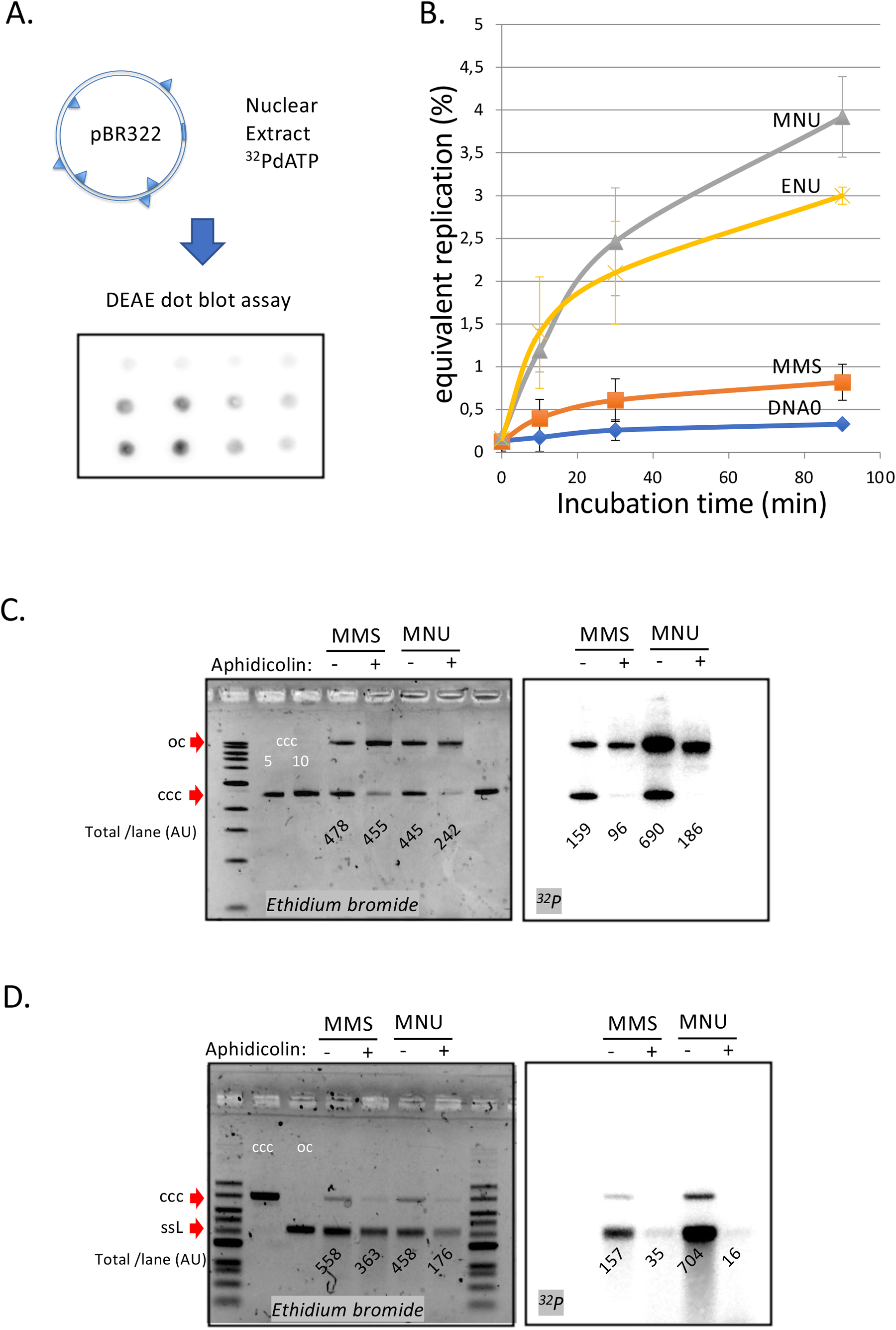
DNA repair synthesis in alkylated and undamaged control plasmid DNA in NPE extracts. **A**. Outline of the spot assay. Plasmids were incubated in nuclear extracts supplemented with α^32^P-dATP; at various time points, an aliquot of the reaction mixture was spotted on DEAE paper (see Material and Methods). **B**. Plasmid DNA pBR322 (4.3kb) samples, modified to a similar extent with -MMS, -MNU and - ENU (see Fig. S1), were incubated in NPE supplemented with α^32^P-dATP at room temperature; incorporation of radioactivity was monitored as a function of time using the spot assay described above (Fig. 1A). Undamaged plasmid DNA0 was used as a control. At each time point, the average values and standard deviation from independent experiments were plotted. The y-axis represents DNA repair synthesis expressed as a fraction of input plasmid replication (i.e. 10% means that the observed extent of repair synthesis is equivalent to 10% of input plasmid replication). This value was determined knowing the average concentration of dATP in the extract (50uM) and the amount of added α^32^P-dATP. **C**. MMS and MNU-treated plasmids were incubated in NPE, supplemented or not, by aphidicolin (150uM final). After 1 h incubation, plasmids were purified and analyzed by agarose gel electrophoresis under neutral loading conditions. The gel was imaged by fluorescence (left: ethidium bromide image) and by autoradiography (right ^32^P image). The number below each lane indicates the total amount of signal per lane (expressed in AU). Aphidicolin treatment decreases incorporation into MNU-treated plasmid close to 4-fold, while it affected incorporation into MMS-treated plasmid only 1.6-fold. **D**. Samples as in C. Gel loading is performed under alkaline conditions to denature DNA before entering the neutral agarose gel allowing single-stranded nick present in DNA to be revealed. The number below each lane indicates the amount of signal per lane (AU).

We asked whether the observed UDS in MNU- and ENU-treated plasmids was MMR dependent, as suggested by the mass spectrometry results. To test this idea, we depleted MMR proteins from extracts using antibodies (Fig. S2A), whose specificity was previously validated (Kato et al., 2017; Kawasoe et al., 2016). Depletion of MLH1 or PMS2 severely reduced UDS in MNU-treated plasmid, while no reduction was observed in PMS1-depleted extracts (Fig. S2A-B). This observation is consistent with the fact that MutLα (composed of MLH1 and PMS2) is involved in canonical mismatch repair whereas MutLβ (composed of MLH1 and PMS1) is not (Jiricny, 2006). Aphidicolin, an inhibitor of B-family DNA polymerases (Baranovskiy et al., 2014), decreased incorporation on average 3.5-fold on MNU and ENU plasmids while it had a more modest effect on MMS-treated plasmid (1.5-fold)(Fig. S2C). These results support the notion that UDS on MNU- and ENU-treated plasmids involves MMR, including a gap filling event that most likely depends on DNA Pol δ, the only B family polymerase detected in the MS analysis described above. Short-patch BER events are mediated by Pol β (X family) that are insensitive to aphidicolin. The modest sensitivity of MMS-induced UDS to aphidicolin is probably due to a fraction of BER events that belong to the long-patch BER pathway mediated Pol δ/*ε* (Sattler et al., 2003).

We wanted to estimate the average amounts of DNA synthesis associated with MMR at O^6^mG:C sites and BER at N-alkyl sites, respectively. At the 90 min time point (i.e. at near plateau value), the difference in UDS between MNU- and MMS-treated plasmids, i.e. attributable to repair at O^6^mG:C sites, was equivalent to ≈ 3.1% of the input DNA (Fig. 2B) or ≈270 nt (pBR322 plasmid is 4363 bp long). With an estimated ≈ 1.7 O^6^mG adducts per plasmid, the average repair patch per O^6^mG adduct is ≈ 160 nt provided all O^6^mG lesions are targeted by MMR. We have evidence that, under the present condition, less than 30% of O^6^mG are substrates for MMR, which suggests that the average MMR patch size is at least 500 nt long. Importantly, the MGMT inhibitor Patrin-2 had no effect on UDS of MNU-treated plasmid, even at a dose of 200 μM (data not shown), suggesting that most O^6^mG adducts do not undergo direct reversal. Surprisingly though, inhibition of MGMT by Patrin-2 was previously shown to occur in Xenopus extracts (Olivera Harris et al., 2015). With respect to N-alkyl adduct repair in MMS-plasmid, repair synthesis above the lesion-free DNA control is equivalent to ≈0.5% of input DNA (Fig. 2B), corresponding to 43 nt total synthesis per plasmid. With ≈ 17 N-alkyl adducts per plasmid, the average DNA synthesis patch per adduct, in case all N-alkyl lesions are repaired, is ≈2.6 nt, a value consistent with a mixture of long (≈ 2-8 nt) and short patch (1nt) BER events at N-alkyl adducts. In summary, the average DNA repair patch sizes at O^6^mG:C (≈500 nt) and N-alkyl (2-3 nt) sites are compatible with MMR and BER, respectively.

To learn more about UDS in this system, we analyzed repair products via gel electrophoresis. Plasmid pBR322 treated with MMS or MNU was incubated in NPE, supplemented or not with Aphidicolin in the presence of α^32^P-dATP, and analyzed on a neutral agarose gel. As already discussed above, addition of Aphidicolin (Aph) led to more severe reduction in incorporation into MNU-(>70%) compared to MMS-treated plasmid (40% reduction) (Fig. 2C). We also note that in MNU-treated plasmids, in the absence of Aph, open circular repair products were three-fold more abundant than closed circular products (Fig. 2C, ^32^P image). This observation suggests that MMR repair was complete in only ≈25% of plasmid molecules while 75% of molecules contained at least one nick (or a gap). Interestingly, there was a ≈50% loss of total DNA in the MNU + Aph lane compared to the other lanes, suggesting massive DNA degradation in NPE due to polymerase inhibition by Aph. Indeed, the observed DNA degradation can specifically be linked to repair events as the loss in radioactivity in MNU lanes -Aph versus + Aph is >70% (Fig. 2C, ^32^P image). Under alkaline loading conditions (Fig. 2D), repair products (^32^P image) in MNU-treated plasmids appeared mostly as a single-stranded linear band form. In addition, there was a large smear (>25% of material) of shorter fragments. These results show that most plasmids contain 1 nick and some contain several nicks. In the +Aph samples, the open circular (oc) form, seen in the gel loaded under neutral conditions (Fig. 2C), disappear under alkali loading conditions (Fig. 2D). This suggests that these oc molecules (Fig 2C) contain many nicks that run as short fragments upon denaturation. In conclusion, MNU-treated plasmids undergo robust repair synthesis that is more sensitive to aphidicolin inhibition than MMS-treated plasmids.

### 3. MMR at single O^6^mG:C base pairs is enhanced by the presence of N-alkylation adducts

Next, we explored a possible crosstalk between repair pathways acting on alkylated DNA. In MNU-treated plasmid, there is on average one O^6^mG adduct for every 9-10 N-alkyl adducts (Beranek, 1990). To investigate the repair response triggered by a single O^6^mG:C lesion alone or in the presence of additional N-alkyl adducts, we implemented a reconstitution experiment. For that purpose, the single O^6^mG:C construct (mGC) (Isogawa et al., 2020) was treated with MMS to introduce ≈ 9-10 N-alkyl adducts per plasmid molecule, generating plasmid mGC+MMS, which is expected to recapitulate adduct distribution found in MNU-treated plasmids. Control plasmid GC was treated with the same concentration of MMS, to generate GC+MMS. These *in vitro* manipulations did not affect plasmid topology as all four constructs exhibit a similar migration pattern (Fig. S3A).

Plasmid constructs GC and mGC and the corresponding two MMS-treated constructs (GC+MMS and mGC+MMS) (Fig. 3A), were incubated with NPE in the presence of α^32^P-dATP to monitor repair synthesis (i.e. UDS). We observed, incorporation of radioactivity specifically attributable to the single O^6^mG:C lesion (compare mGC to GC in Fig. 3B). Activation of MMR by a single O^6^mG:C lesion has been reported previously (Duckett et al., 1999). The specific involvement of MMR for O^6^mG dependent incorporation was re-assessed, by incubating the single adducted O^6^mG:C construct in MLH1-depleted NPE extract; radioactive incorporation above background was fully abolished in mGC plasmid (Fig. S3E). How MMR may get engaged in a repair reaction on a closed circular template will be discussed in the Discussion section.

**Fig. 3.**
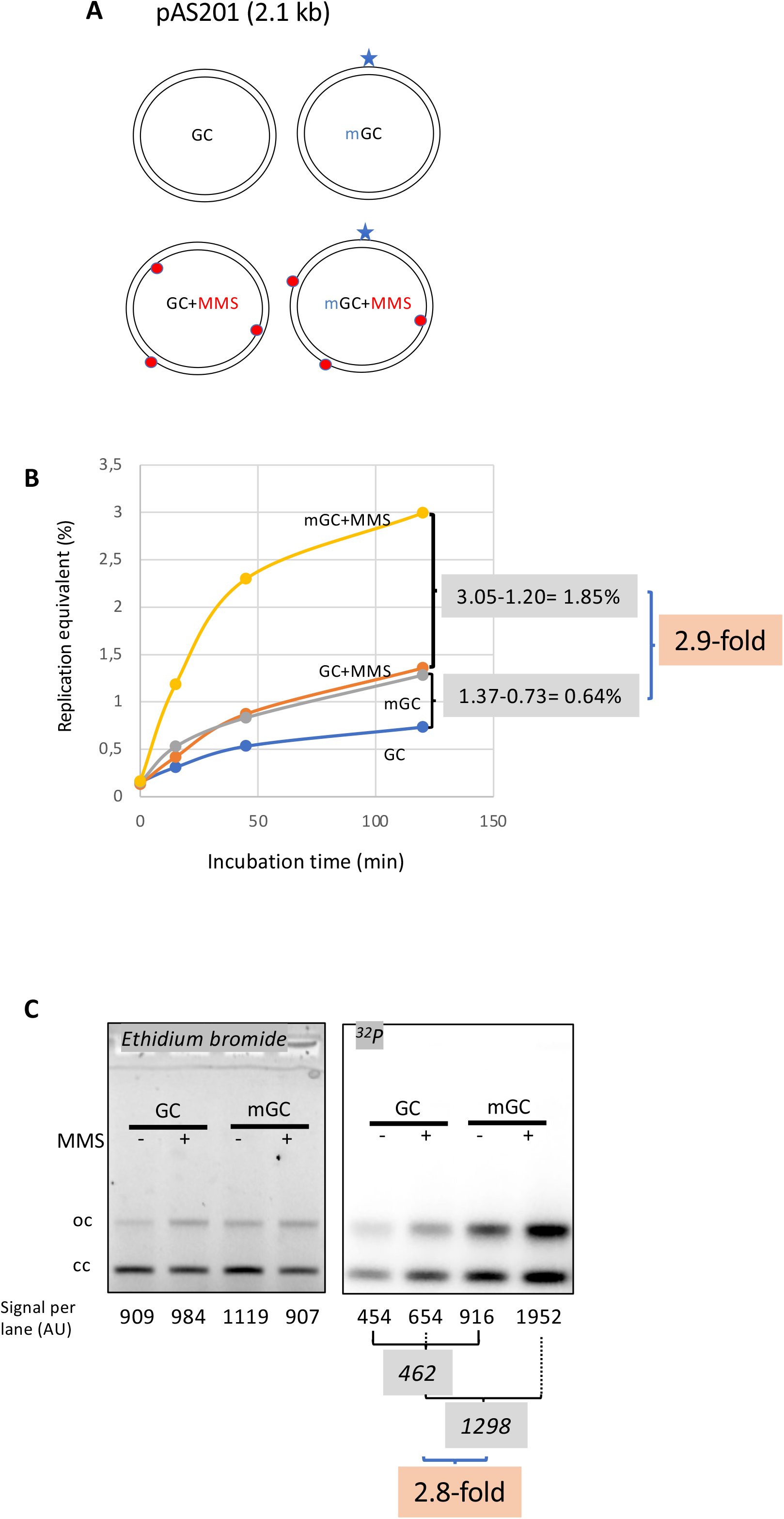
Stimulation of MMR at a single O^6^mG site by N-alkyl adducts *in cis*. **A**. Covalently closed circular (ccc) plasmids (pAS201, 2,1 kb) containing a site-specific O^6^mG:C base pair (plasmid mGC) and the corresponding lesion-free control (plasmid GC) were constructed (Isogawa et al., 2020). Both GC and mGC plasmids were treated with MMS in order to introduce random N-alkyl (7mG and 3mA) adducts, generating plasmid GC+MMS and mGC+MMS, respectively. We adjusted the MMS reaction conditions as to introduce ≈ 9 adducts per plasmid (i.e. one N-alkylation adduct every 4 to 500 nt). The resulting proportion of O-alk and N-alkyl adducts mimics the proportion in MNU-treated plasmid. The single O^6^mG adduct and the randomly located N-alkyl adducts are represented by a star and red dots, respectively. **B**. Plasmids described above were incubated in NPE supplemented with α^32^P-dATP at room temperature; incorporation of radioactivity was monitored as a function of time using the spot assay described above (Fig. 2A). The y-axis represents the percentage of DNA repair synthesis with respect to input DNA (i.e. 100% would mean that all input DNA had been newly synthesized: as in 2B). Incorporation attributable to repair at the O^6^mG:C lesion is increased close to 3-fold due to the presence of random N-alkyl lesions introduced by MMS treatment. **C**. The same plasmids were incubated for 2h incubation in NPE, purified, resolved by agarose gel electrophoresis and revealed by ethidium bromide fluorescence and ^32^P autoradiography. The total amount of signal per lane is indicated (AU). As expected, the amount of plasmid extracted from each incubation mix is relatively constant, as quantified below the ethidium bromide image. Increase in repair at the O^6^mG:C lesion due to MMS-treatment (2.8-fold) is in good agreement with data in 3B.

Importantly, repair synthesis, due to the single O^6^mG:C lesion, is strongly enhanced by the presence of MMS adducts (compare mGC+MMS to GC+MMS in Fig. 3B). At the 2h time point, incorporation, above background, due to the single O^6^mG, expressed in % replication equivalent, represents 0.64% (difference between mGC and GC), while it amounts to 1.85% in the presence of MMS lesions (compare mGC+MMS with GC+MMS). One can thus estimate that, incorporation due to a single O^6^mG lesion, is stimulated about 2.9-fold (1.85/0.64) by the presence in *cis* of MMS adducts (Fig 3B).

### 4. Nucleotide incorporation occurs in the vicinity of the single O^6^mG adduct

The plasmids described above were incubated in α^32^P-dATP supplemented NPE for 2h, purified plasmids and analyzed by agarose gel electrophoresis (Fig. 3C). Covalently closed circular (ccc) and relaxed forms (oc) were quantified in each lane (as indicated in Fig. 3C). In the presence of MMS adducts, the single O^6^mG:C lesion contributes to a 2.8 fold increase in radioactive incorporation compared to its contribution in the absence of MMS (Fig 3C) in good agreement with the UDS data (Fig. 3B).

We wanted to map the repair patches with respect to the position of the O^6^mG adduct by restriction enzyme analysis by digesting the purified plasmids with *BmtI* and *BaeGI* restriction enzymes, generating fragment S (589 bp) that encompasses the O^6^mG:C site and fragment L (1525 bp) (Fig. S3B). Following separation by agarose gel electrophoresis, the DNA was imaged by ethidium bromide fluorescence and ^32^P autoradiography (Fig. S3C). For each fragment, we determined its specific activity by dividing its radioactivity with its total abundance determined from the ethidium bromide image (Fig. S3D). As expected, the specific activities of S and L fragments were similar in GC control plasmid (random background incorporation: 0.125±0.015 AU (arbitrary units)) and MMS treated (GC+MMS) (0.235±0.025 AU) control plasmid. In GC+MMS, the specific activity was slightly higher than in control plasmid, probably reflecting BER-mediated incorporation at randomly distributed N-alkyl adducts. In the two O^6^mG:C containing plasmids (mGC and mGC+MMS), the S fragment exhibits a significantly higher specific activity than the L fragment indicating that MMR activity is centered around the O^6^mG:C site. In the absence of MMS, incorporation in mGC above background (dotted line in S3D), attributable to O^6^mG, amounts to 0.065 and 0.495 AU (arbitrary units) for L and S fragments, respectively. Similarly, in the presence of random MMS lesions (mGC+MMS), incorporation, above background (dotted line in S3D), attributable to O^6^mG amounts to 0.115 and 1,17 AU for L and S fragments, respectively. These results clearly show that O^6^mGC induced repair essentially takes place within the S fragment, with only modest spill-over into the L fragment (10-15%). This observation appears to be good agreement with the estimated average MMR patch size (∼500 nt). Thus, MMS adducts do not modify the repair pattern, i.e. the relative distribution of ^32^P incorporation in S and L fragments, but they increase the frequency of repair centered at O^6^mG sites. In conclusion, we show that stimulation of repair synthesis by N-alkyl adducts specifically occurs in the vicinity of the O^6^mG adducts, illustrating that processing of N-alkyl adducts enhances MMR activity.

We next examined O^6^mG-induced DNA synthesis in a different extract, High Speed Supernatant (HSS) of total egg lysate. Although HSS is proficient for MMR at a single O^6^mG flanked by a nick (Olivera Harris et al., 2015), HSS contains lower concentrations of most DNA repair enzymes compared to NPE. Interestingly, in contrast to NPE, HSS did not support incorporation of α^32^P-dATP into MNU-treated plasmids (Fig. S4). We reasoned that HSS might not contain adequate concentrations of the DNA glycosylase AAG, which initiates BER at N-alkyl sites. Indeed, in HSS supplemented with human AAG, the UDS response triggered in MNU-treated plasmid was well above that observed on MMS-treated plasmid (Fig. S4). These observations support the involvement of BER in stimulating MMR at O^6^mG lesions.

### 5. MNU-treated plasmids undergo double-strand breaks during incubation in extracts

Work in *E. coli*, provided elegant genetic evidence that the cytotoxicity of alkylating agents forming O^6^mG adducts (such as N-methyl-N’-nitrosoguanidine and MNU), including formation of replication-independent DSB, was strongly influenced by the status of the MMR pathway (Karran and Marinus, 1982; Nowosielska and Marinus, 2008). We wondered whether MNU can induce formation of DSBs independently of DNA replication. To increase the sensitivity of our assay towards DSB detection, we used a larger plasmid, pEL97 (11,3 kb), and treated it with MMS or MNU to introduce one alkylation event on average every 500 nt (Fig. S5). We also treated one sample with double the concentration of MNU (MNU++, 1 lesion every 250 nt). Quantification of N-alkyl adducts by alkaline cleavage (Fig. S1A) and subsequent agarose gel electrophoresis led to the expected lesion density (Fig. S5C). The extent of alkylation triggered by MNU as deduced from the alkaline cleavage determination fits surprisingly well with the amount of alkylation induced by TMZ in cell (Kaina, personal communication). Alkylated and control plasmids (Fig. S6A) were incubated in NPE for 60’ in the presence of α^32^P-dATP, resolved by agarose gel electrophoresis, and visualized by ethidium bromide staining and ^32^P imaging (Fig. 4A). Both MMS and MNU caused substantial conversion of the plasmid from supercoiled to open circular form, as expected during repair synthesis. Consistent with our results above, MNU induced much more repair synthesis than MMS. Strikingly, in both ethidium bromide and ^32^P images, linear plasmid was detected after exposure to MNU, but not MMS. For a two-fold increase in MNU exposure, the linear plasmid band increased ≈4-fold (Fig. 4B). The quadratic dose-response (Fig. 4B) strongly suggests that DSBs occur as a consequence of two independent repair events at neighboring lesions, for example a BER event at an N-alkyl adduct leading to a nick in one strand that is encountered by a gap formed by an MMR event initiated at a O^6^mG site in the opposite strand (Fig. 5). To reveal all single-strand breaks, the same samples were also denatured prior to native gel electrophoresis (Fig. S6B). In the MNU++ sample, no linear single-stranded DNA (ssDNA) is left, with the DNA molecules running as a smear around the 3000 nt position (Fig. S6B). The observed smear reveals that the double-stranded DNA running as open circular plasmid molecules in the neutral loading gel (Fig. 4A), contain each on average 3-4 nicks per plasmid strand. The data reveal that repair of MNU-treated plasmid in Xenopus egg extracts causes SSBs, and that once the density of SSBs is high enough, DSBs result.

**Fig. 4.**
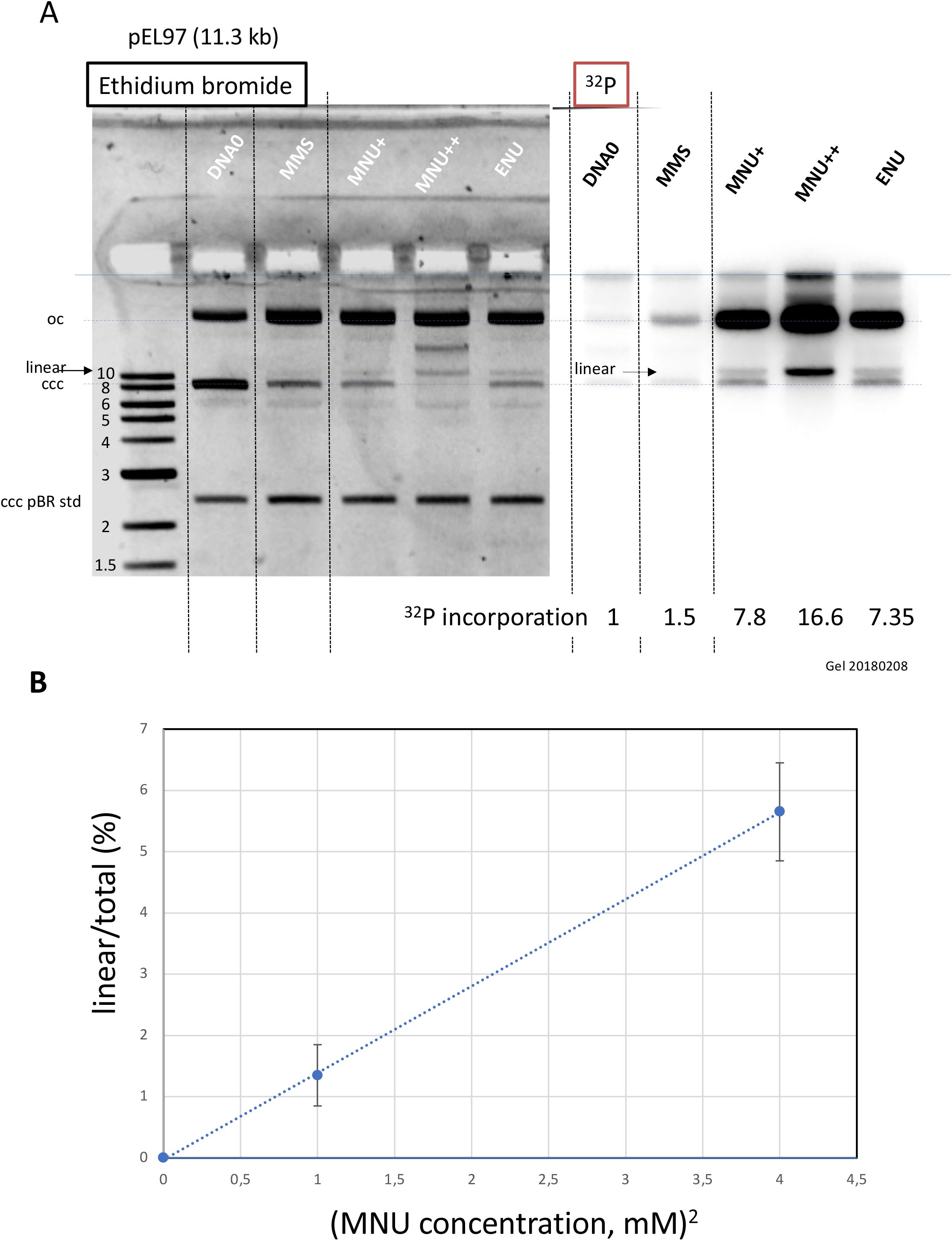
Double-strand breaks occur in MNU-treated plasmid during incubation in extracts. **A**. Analysis by AGE of alkylated plasmids (pEL97: 11,3kb) incubated in NPE in the presence of α^32^P-dATP. Plasmid pEL97 was treated with MMS, MNU+ and ENU as to introduce ≈ one alkylation event on average every 500 nt on average. For MNU, a plasmid with twice the level of alkylation (MNU++, 1 lesion every 250 nt) was also produced (Fig. S5). Alkylation of these plasmids essentially not affected their migration on agarose gels (Fig. S6A). After 2h incubation, the reaction was stopped and a known amount of pBR322 (10 ng) plasmid was added as an internal standard. After electrophoresis, the gel was imaged by fluorescence (ethidium bromide staining) and by autoradiography. *Ethidium bromide image*: in the different lanes the internal standard band, pBR (ccc), appears to be of similar intensity (1158 +/-95 AU), assessing reproducible DNA extraction. For the alkylated plasmids, incubation in NPE led to massive conversion from ccc to relaxed plasmid. ^*32*^*P image*: Little incorporation of ^32^P-dATP is seen in DNA0 and in MMS-treated plasmid compared to MNU and ENU treated plasmids as shown by the relative incorporation levels normalized to 1 for untreated plasmid (DNA0). As expected, the MNU++ sample exhibits about twice the amount of incorporated radioactivity compared to MNU+ (see relative amount of incorporated radioactivity). In both Ethidium bromide and ^32^P images, a small amount of linear plasmid is seen mostly in the MNU++ sample. This band is also visible in the MNU+ and ENU lanes although at a weaker intensity. **B**. Quadratic dose-response for DSB formation. When the % of linear form (linear/(linear + oc)), is plotted as a function of the square dose of MNU (mM^2^) for untreated, MNU+ and MNU++ plasmids, we observed a straight line (y = 1,4173x - 0,0288; R^2^ = 0,9999).

**Fig. 5:**
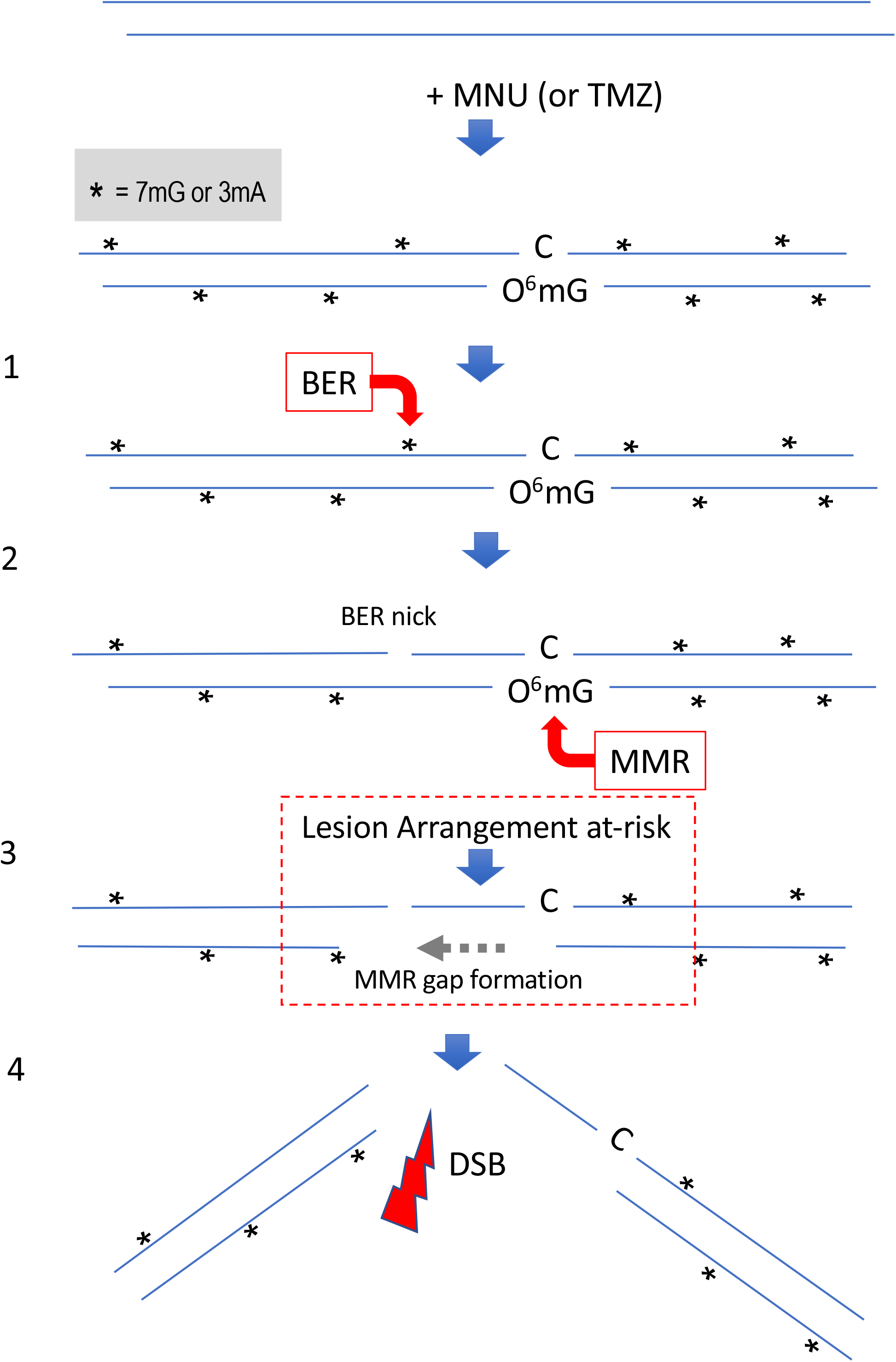
Simultaneous repair of two closely spaced MNU-induced lesions may lead to a DSB. Such a situation occurs when an N-alkyl lesion located within ≈1000nt of an O^6^mG lesion are processed simultaneously (“Lesion Arrangement at-risk”). Note that the MMR excision track can occur on the opposite strand as described for noncanonical MMR (Peña-Diaz et al., 2012). Reaction of MNU with double-stranded DNA induces N-alkylation adducts, mostly 7mG and 3mA shown as * and O-alkylation adducts (O^6^mG), at a 10:1 ratio approximatively. Step 1: a BER event is initiated at an N-alkyl adduct, creating a nick. Step 2: concomitantly, a MMR event takes place, in the opposite strand, at a nearby at a O^6^mG:C site. Step 3: the MMR machinery extends the nick into a several hundred nt long gap by means of Exo1 action. Step 4: The two independently initiated repair events lead to a DSB, if the MMR gap reaches the BER initiated nick before resealing.

## Discussion

With respect to the biological responses to SN1 alkylating agents most attention has so far been devoted to responses that occur in the first or second cell cycle following treatment as mentioned in the introduction (Noonan et al., 2012; Plant and Roberts, 1971; Quiros et al., 2010).

In the present paper, we focus on early processing of DNA alkylation adducts by repair pathways independently of replication. Interestingly, we identify the formation of DSB as the result of a putative crosstalk between repair pathways.

### Late responses to SN1 agents

Response of cells to SN1 methylating agents was shown to be initiated at O^6^mG:T mispairs that form upon DNA replication of O^6^mG containing DNA template and shown to involve the MMR machinery (Goldmacher et al., 1986) (Day et al., 1980; Karran et al., 1993; Yarosh et al., 1983). The O^6^mG:T mispair is efficiently recognized by MutSα, the key MMR initiator protein. Following removal of the nascent T residue across O^6^mG, T will be re-inserted during the MMR gap-filling step thus re-forming the initial O^6^mG:T mispair. This iterative process, called “futile cycling”, has received experimental support (Mazon et al., 2010; York and Modrich, 2006). However, it is not yet clear how these futile cycles lead to DSBs (Ochs and Kaina, 2000), apoptosis and cell death (Gupta and Heinen, 2019; Kaina and Christmann, 2019). Two mutually non-exclusive models have been proposed, i) a direct model where the encounter of the replication fork with the MMR intermediates leads to fork collapse and to subsequent cytotoxic events; ii) a signaling model where the MutSα complex acts as a sensor leading to ATR recruitment and subsequent initiation of the ATR-Chk1 signaling pathway (Duckett et al., 1999; Yoshioka et al., 2006). However, presently there is more evidence that the critical cytotoxic response to methylating agents is the consequence of direct MMR processing rather than being mediated by a mere signaling model (Cejka and Jiricny, 2008; Karran, 2001; Liu et al., 2010; York and Modrich, 2006).

### Early responses to SN1 agents

While all biological responses described above require replication of O^6^mG containing DNA templates as the first step, we wanted to investigate the processing of MNU-alkylated DNA in the absence of replication. Interestingly, we detected robust UDS upon incubation of MNU-treated plasmid in Xenopus egg extracts that was shown to be dependent upon MMR proteins. This observation indicated that, not only is the O^6^mG:C lesion recognized by MutSα as previously noted (Duckett et al., 1999; Karran et al., 1993), but that the whole repair process proceeds to completion.

Agents such as MNU and TMZ induce about 10 times more N-alkyl adducts compared to O^6^mG. We wanted to investigate the potential effect that N-alkyl adducts may have on MMR processing at O^6^mG:C base pairs. For that purpose, we compared a plasmid with a single site-specific O^6^mG:C lesion to a plasmid additionally treated with MMS, an agent known to induce essentially only N-alkyl adducts. The MMS treatment was adjusted as to produce the same amount of N-alkyl adduct as generated by MNU. The single-adducted O^6^mG:C plasmid triggered MMR mediated repair synthesis centered around the O^6^mG adduct. Interestingly, the presence of randomly distributed N-alkyl adducts led to a 3-fold increase of the MMR repair activity in the vicinity of the O^6^mG adduct. These data raise two questions, first, how does the MMR machinery get engaged in ccc plasmid and how is MMR activity further stimulated by N-alkyl adducts? In current models, functional engagement of MMR involves a mismatch recognized by MutSα, subsequent recruitment of MutLα and PCNA (Jiricny, 2006). Loading of PCNA by RFC normally requires a single-stranded nick but it was also shown to occur, although less efficiently, on ccc DNA (Pluciennik et al., 2013; 2010). Under these conditions, PCNA loading and MMR processing lack strand directionality. With respect to the mechanism by which MMR activity, at a single O^6^mG:C lesion, becomes stimulated several folds by the presence of N-alkyl adducts, we propose that processing of N-alkyl adducts by BER creates repair intermediates (nicks) that stimulate PCNA loading. It was previously shown that BER intermediates formed at oxidized purines or U residues can stimulate MMR processing (Repmann et al., 2015; Schanz et al., 2009).

### DSB form in MNU-treated DNA in the absence of replication: potential therapeutic significance

Interestingly, incubation of MNU-treated plasmid in extracts leads to DSBs (Fig. 4) that arise with a quadratic dose response suggesting crosstalk between two independent repair activities taking place simultaneously in opposite strands at lesions that may be up to several hundred nucleotides apart (see scenario in Fig. 5). Similarly, *in vitro* processing of neighboring G/U mispairs by BER and by non-canonical MMR was shown to lead to DSBs (Bregenhorn et al., 2016).

According to the model (Fig. 5), formation of a DSB may occur when an N-alkyl lesion is located within the repair track mediated by MMR at a O^6^mG:C site. We can estimate the number of events (per genome) where an N-alky lesion is located within 500 nt on either side of a O^6^mG:C site. In the clinic, a daily dose of TMZ results in 50uM serum concentration that was shown to induce 5.2×10^4^ and 7.3×10^5^ O-alkyl and N-alkyl lesions per human genome, respectively (Kaina, personal communication). Given the N-alkyl lesion density (7.3×10^5^/6×10^9^= 1.2×10^−4^), the probability of presence of an N-alky lesion within an MMR track is 0.12. In other words, among the 5.2×10^4^ O^6^mG lesions, ≈6,240 are likely to contains a N-alkyl lesion within a 1000 nt excision track. We will refer to such a lesion configuration as a “Lesion Arrangement at-risk” for DSB formation.

Let us now estimate the level of DSB that may occur in a human genome, by extrapolation. In the present work, ≈6% of plasmid DNA (11.3 kb) treated by MNU at 2mM exhibit a DSB (Fig. 4). The observed amount of DSBs may in fact only reflect a steady state level since efficient re-ligation mechanisms are known to operate in Xenopus egg extracts (Graham et al., 2016). Since MNU and TMZ exhibit similar reactivities (Moody and Wheelhouse, 2014), a dose of 2mM MNU would lead to 3×10^9^ x 0.06 / 11300= 16,000 DSBs per genome. In the clinic, the measured level of TMZ in the serum reaches up to 50uM, i.e. 40 times less than the 2mM dose used here. Given the quadratic dose-response, the estimated amount of DSBs per genome would be 1600 times less, i.e. ≈10. Lets note that the conversion of a Lesion Arrangement at-risk into an actual DSB appears to be quite low (10/6,240 ≈0,16%), reflecting the requirement for simultaneous occurrence of the two repair events (MMR and BER). While 10 DSB/cell is already a significant amount, it should be kept in mind that under clinical regimens TMZ treatment occurs daily for weeks thus likely leading to daily formation and accumulation of this class of DSBs.

Thus, the number of DSBs/cell induced by TMZ, in the absence of replication, as a result of the crosstalk between MMR and BER is comparable to the number of DSBs induced by 0.5-1 Gy of IR. The alkylating agent Temozolomide (TMZ), a chemical mimic of MNU, is presently the first-line and only anti-cancer drug in glioblastoma therapy (Moody and Wheelhouse, 2014). Understanding both early and late cellular responses to MNU/TMZ appears thus to be critical. During cancer treatment, a dose of TMZ is delivered concomitantly with a radiotherapy session daily, for 6 weeks (for a recent review see Strobel et al., 2019). Moreover, it was established empirically, that the treatment TMZ plus radiotherapy exhibits supra-additive cytotoxicity as long as TMZ administration *precedes* radiotherapy (Bobola et al., 2010). Our data may provide some rational for this empirically determined regimen. Indeed, the single-stranded DNA stretches formed at early time points during MMR processing at O^6^mG:C sites (step 3 in Fig. 5) represent preferential targets for conversion of the numerous single-stranded breaks induced by ionizing radiations (IR) into DSBs, thus providing an explanation for the observed supra-additivity in the treatment when TMZ *precedes* IR. A commonly used radiotherapy session involves an IR dose of 2 Gy that predominantly induces ≈2000 single-strand breaks and 40-80 DSBs/cell.

In conclusion, the present work offers a novel mechanistic insight into the cytotoxicity of TMZ via induction of DSBs, at early time points following exposure, before replication. This early response comes in complement to the late, replication and cell cycle dependent, responses that have been described over the years. This additional mechanism for TMZ cytotoxicity will need to be investigated in cellular systems.

## Acknowledgements

We thank Johannes Walter (Harvard Medical School) for providing space, advice and materials. Paul Modrich (Duke Univ) and Bernd Kaina (Medical Univ, Mainz) for insightful reading and suggestions.

## Material and Methods

### Akylation reaction

are conducted as indicated in the table below, at a plasmid concentration of 10ng/ul in CE buffer (citrate 10mM pH7,EDTA 1mM) + 10% DMSO final.

| 0.65 ml tubes | DNA0 | MMS: 10mM<br>final | MNU: 1mM<br>final | ENU: 30mM<br>final |
| --- | --- | --- | --- | --- |
| pBR322 1ug in CE | 90ul | 90ul | 90ul | 90ul |
| Alkylating agent in DMSO | 10ul DMSO | 10ul MMS at 100mM | 10ul MNU at 10mM | 10ul ENU at 300mM |
| Temp / time | 50°C / 40 min | 40°C / 30min | 50°C / 15-17 min | 50°C / 40 min |

The alkylation reactions are terminated by addition of STOP buffer (5x: 1.5M sodium acetate, 1M mercapto-ethanol) followed by ethanol precipitation. The DNA pellet is washed with ethanol 90% and re-dissolved in TE at 50ng/ul (see native agarose gel profile Fig. S1A).

### Cleavage reactions at 7-alkylG and 3-alkylA adducts

Alkylated plasmids (50ng in 10ul of CE buffer) are first incubated for 90°C during 15’ at pH7 (PCR machine). Following addition of 1ul of NaOH 1N, the sample is further incubated at 90°C for 30’. Following addition of 2ul of alkaline 6x loading buffer (NaOH 300mM, EDTA 6mM, Ficoll (Pharmacia type 400) 180mg/ml, 0.15% (w/v) bromocresol green, 0.25% (w/v) xylene cyanol), the cleaved plasmid samples are loaded on a neutral agarose gel.

### NPE and HSS Xenopus extracts

Preparation of *Xenopus* egg extracts was performed as described previously (Lebofsky et al., 2009).

### Western Blot

Antibodies against Mlh1, Pms2 and Pms1 as previously described (Kato et al., 2017; Kawasoe et al., 2016). For western blotting primary antibodies used at 1:5000 dilution.

### Plasmid immobilization on magnetic beads

Covalently closed circular plasmids (pAS201, 2,1 kb) containing a site-specific O^6^mG:C base pair (plasmid mGC) and the corresponding lesion-free control (plasmid GC) were derived from plasmid pAS200.2 (Isogawa et al., 2020). Alkylated plasmids samples (250 ng of each -MMS, - MNU and -ENU), as well as a non-alkylated control sample (DNA0) were immobilized on magnetic beads at a density of 10 ng plasmid / ul of M280 bead slurry using a triple helix-based capture methodology (Isogawa et al., 2018). The TFO1 probe used here is 5’ Psoralen – C6 – TTTT<u>C</u>TTTT<u>C</u>T<u>CC</u>T<u>C</u>TT<u>C</u>T<u>C</u>– C124 –Desthiobiotin (20 mer) with C124: hexaethylene glycol ×6. Underlined C is for 5mC; it was synthesized by using DNA/RNA automated synthesizer and purified with conventional methods (Nagatsugi et al., 2003).

In order to monitor non-specific protein binding to beads, we included an additional control with the same amount of M280 beads without plasmid DNA (noDNA).

Immobilized plasmid DNA are incubated in Xenopus egg extract (NPE) for 10 min at room temperature under mild agitation. The reaction was stopped by addition of 320ul of a 0.8% HCHO solution to cross-link the proteins-DNA complexes for 10 min at room temperature. The beads were subsequently washed at RT with 200ul of 100mM NaCl containing buffer (ELB buffer), re-suspended in 70ul of Extract Dilution buffer and layered on top of a 0.5M sucrose cushion in a long Beckman centrifugation tube. The beads were quickly spun trough the cushion (30 sec at 10 000 rpm), the bead pellet is re-suspended into 40ul of ELB sucrose and further analyzed by PAGE or by MS.

### Incorporation of α^32^P-dATP into DNA: spot assay

Plasmids are incubated in nuclear extracts supplemented with α^32^P-dATP; at various time points, an aliquot of the reaction mixture is spotted on DEAE paper (DE81). The paper is soaked in 100ml 0.5M Na2HPO4 (pH≈9), shacked gently for 5’ before the buffer is discarded; this procedure is repeated twice. Finally, the paper is washed an additional 2 times in 50ml ethanol, air dried and analyzed by ^32^P imaging and quantification. The extent of DNA repair synthesis is expressed as a fraction of input plasmid replication (i.e. 10% means that the observed extent of repair synthesis is equivalent to 10% of input plasmid replication). This value is determined knowing the average concentration of dATP in the extracts (≈50uM) and the amount of added α^32^P-dATP.

### PAGE / silver staining

An aliquot of each incubation experiments, corresponding to 30 ng of immobilized plasmid was treated at 99°C for 25 min in a PCR machine to revert HCHO cross-linking in LLB, 50 mM DTT. Samples were loaded on a 4-15% PAGE (Biorad pre-cast) gel at 200 volts for 32min and stained using the silver staining kit (silver StainPlus, Biorad).

### Mass Spectrometry

Label-free mass spectrometry analysis was performed using on-bead digestion. In solution digestion was performed on beads from plasmid pull-downs. We added 20 µl of 8 M urea, 100 mM EPPS pH 8.5 to the beads, then 5mM TCEP and incubated the mixture for 15 min at room temperature. We then added 10 mM of iodoacetamide for 15min at room temperature in the dark. We added 15 mM DTT to consume any unreacted iodoacetamide. We added 180µl of 100 mM EPPS pH 8.5. to reduce the urea concentration to <1 M, 1 µg of trypsin, and incubated at 37°C for 6 hrs. The solution was acidified with 2% formic acid and the digested peptides were desalted via StageTip, dried via vacuum centrifugation, and reconstituted in 5% acetonitrile, 5% formic acid for LC-MS/MS processing.

All label-free mass spectrometry data were collected using a Q Exactive mass spectrometer (Thermo Fisher Scientific, San Jose, CA) coupled with a Famos Autosampler (LC Packings) and an Accela600 liquid chromatography (LC) pump (Thermo Fisher Scientific). Peptides were separated on a 100 μm inner diameter microcapillary column packed with ∼20 cm of Accucore C18 resin (2.6 μm, 150 Å, Thermo Fisher Scientific). For each analysis, we loaded ∼2 μg onto the column. Peptides were separated using a 1 hr gradient of 5 to 29% acetonitrile in 0.125% formic acid with a flow rate of ∼300 nL/min. The scan sequence began with an Orbitrap MS^1^ spectrum with the following parameters: resolution 70,000, scan range 300-1500 Th, automatic gain control (AGC) target 1 × 10^5^, maximum injection time 250 ms, and centroid spectrum data type. We selected the top twenty precursors for MS^2^ analysis which consisted of HCD high-energy collision dissociation with the following parameters: resolution 17,500, AGC 1 × 10^5^, maximum injection time 60 ms, isolation window 2 Th, normalized collision energy (NCE) 25, and centroid spectrum data type. The underfill ratio was set at 9%, which corresponds to a 1.5 × 10^5^ intensity threshold. In addition, unassigned and singly charged species were excluded from MS^2^ analysis and dynamic exclusion was set to automatic.

### Mass spectrometric data analysis

Mass spectra were processed using a Sequest-based in-house software pipeline. MS spectra were converted to mzXML using a modified version of ReAdW.exe. Database searching included all entries from the *Xenopus laevis*, which was concatenated with a reverse database composed of all protein sequences in reversed order. Searches were performed using a 50 ppm precursor ion tolerance. Product ion tolerance was set to 0.03 Th. Carbamidomethylation of cysteine residues (+57.0215Da) were set as static modifications, while oxidation of methionine residues (+15.9949 Da) was set as a variable modification. Peptide spectral matches (PSMs) were altered to a 1% FDR (Elias and Gygi, 2010; 2007). PSM filtering was performed using a linear discriminant analysis, as described previously (Huttlin et al., 2017), while considering the following parameters: XCorr, ΔCn, missed cleavages, peptide length, charge state, and precursor mass accuracy. Peptide-spectral matches were identified, quantified, and collapsed to a 1% FDR and then further collapsed to a final protein-level FDR of 1%. Furthermore, protein assembly was guided by principles of parsimony to produce the smallest set of proteins necessary to account for all observed peptides.

## Supplementary information

### Reaction conditions leading to similar levels of DNA adduct formation

As a proxy for total alkylation we decided to monitor the two major N-alkyl adducts, namely 7mG and 3mA, that together represent > 80% of alkylation for MMS and MNU. These N-alkyl adducts were converted into ss DNA breaks by a combination of heat depurination and alkali cleavage treatment (Maxam and Gilbert, 1977)(see Fig. S1A) and the resulting plasmid fragmentation pattern were analyzed by agarose gel electrophoresis. The reaction conditions were adjusted as to generate a median fragment size of 500 nt, corresponding to one alkylated base every 500 nucleotides on average (Fig S1A).

## Legend to supplementary figures

**Fig. S1.**
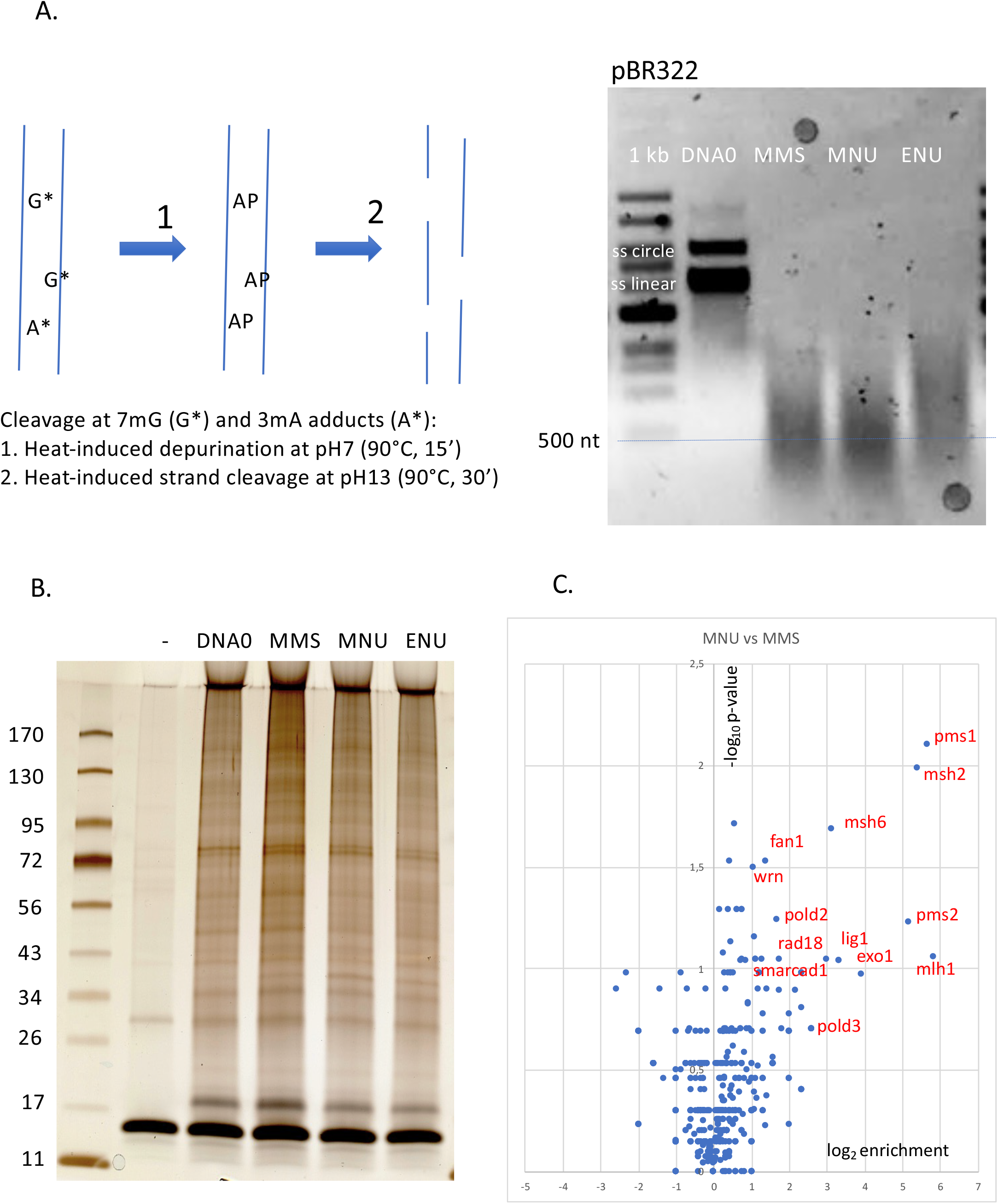
<u>A. Alkaline cleavage procedure at N-alkylation adducts (7mG and 3mA)</u>. Alkylated pBR322 plasmids (50ng in 10ul of CE buffer) were first incubated for 90°C during 15’ at pH7 (PCR machine). Following addition of 1ul of NaOH 1N, the sample was further incubated at 90°C for 30’. Following addition of 2ul of alkaline 6x loading buffer (NaOH 300mM, EDTA 6mM, Ficoll (Pharmacia type 400) 180mg/ml, 0.15% (w/v) bromocresol green, 0.25% (w/v) xylene cyanol) the cleaved plasmid samples were resolved by neutral agarose gel electrophoresis. <u>B. Plasmid pull-down: protein analysis by PAGE and silver staining</u>: Plasmid, pAS04 (6,47 kb) was modified by MMS, MNU and ENU to a similar extent corresponding to an average of one N-alkyl adduct every 500 nt as described in Fig. S1A. Control plasmid (DNA0) and alkylated plasmids were incubated in NPE as outlined in Fig. 1A for 10min at room temperature. Samples, corresponding to ≈ 30 ng of input plasmid DNA, were loaded on a 4-15% PAGE (Biorad pre-cast) gel, run at 200 volts for 32min and silver stained. Lanes DNA0, MMS, MNU and ENU display a complex pattern of proteins (total amount of proteins per lane estimated at 100-200 ng). Interestingly, the noDNA lane shows a low amount of residual proteins indicating the efficiency of the post-incubation bead isolation procedure to remove unspecific proteins. <u>C. Relative abundance of proteins captured on MNU-versus MMS-treated plasmid</u>. Proteins captured on equal amounts of MNU- or MMS-treated plasmid were analyzed by label-free MS in triplicates. The data are analyzed and plotted as described in Fig. 1B using Xenbase data base. Proteins enriched on MNU versus MMS-treated DNA appear in the right-side top corner and essentially turn out to be MMR proteins labelled in red (Fig. 1B).

**Fig. S2.**
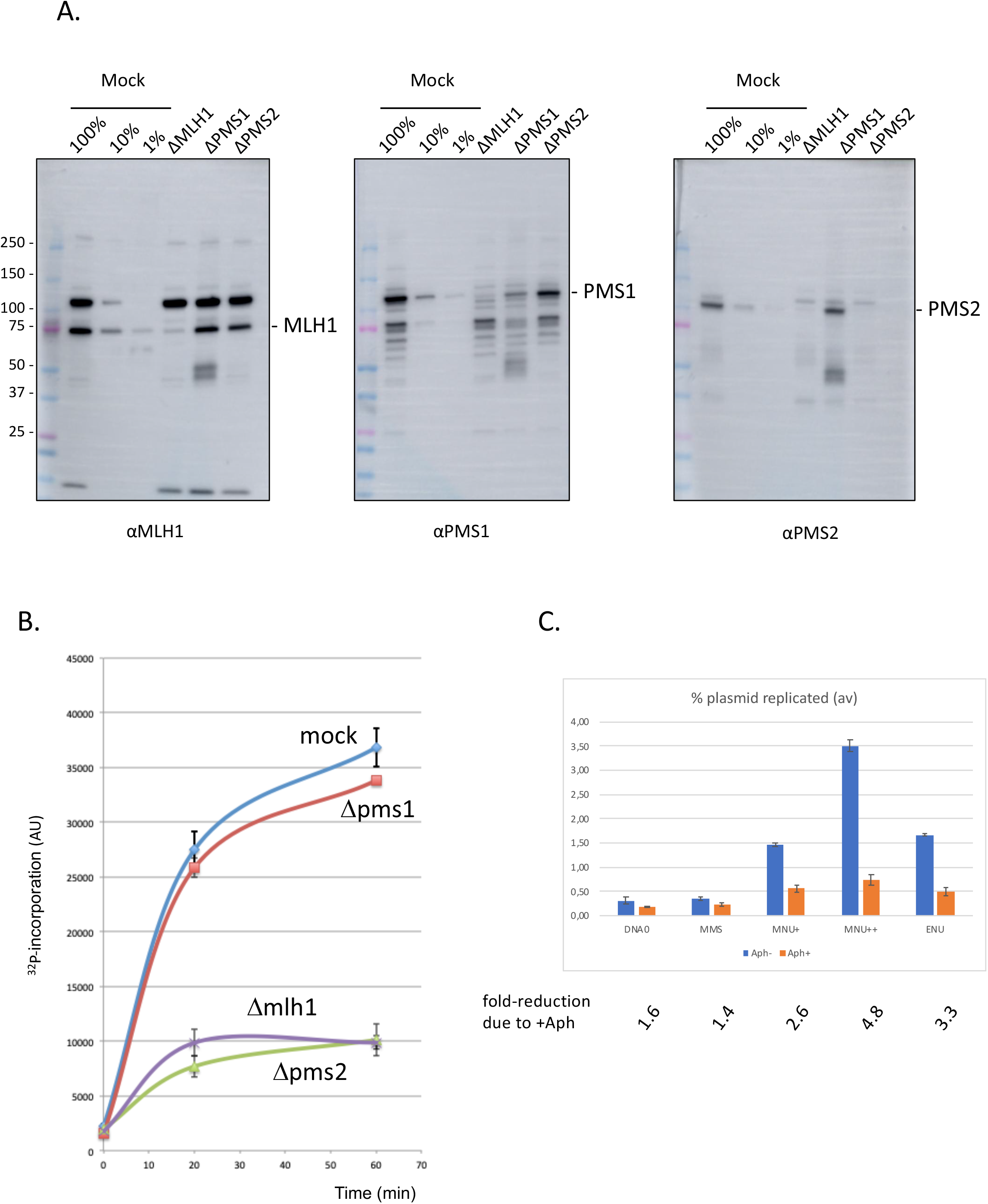
Involvement of Mismatch repair in repair synthesis and effect of aphidicolin. **A**. Analysis by WB of the NPE extracts depleted for MMR proteins. Antibodies against Mlh1, Pms2 and Pms1 as previously described (Kato et al., 2017; Kawasoe et al., 2016). Extracts were depleted, with the respective antibodies, for three rounds at 4C° at a ratio beads: antibody: extract = 1 :3 :5. **B:** Depletion experiments show that α^32^P-dATP incorporation into MNU-plasmid is mediated by Mismatch Repair. NPE was depleted by antibodies against Pms1, Pms2, Mlh1 or mock depleted. Upon incubation at room temperature in the different NPE extracts, incorporation of α^32^P-dATP in MNU-plasmid was monitored by the spot assay as a function of time. At each time point, the average values and standard deviations from two independent experiments were plotted. As expected from previous data (Fig. 2B), robust incorporation was observed for MNU-plasmid incubated in mock-depleted extract. In contrast, radioactive dATP incorporation was severely reduced when the plasmid was incubated in extracts depleted with antibodies against Mlh1 or Pms2. These data strongly suggest that the incorporation seen in mock depleted NPE is mediated by MMR, as both Mlh1 and Pms2 assemble into the functional MMR complex MutLα. In contrast, depletion with antibodies against Pms1 did not affect incorporation kinetics as Pms1 is reported not to function in MMR (Jiricny, 2006). **C**. Effect of aphidicolin: alkylated plasmids (pEL97: 11.3kb) were incubated in NPE supplemented of not by aphidicolin (150uM final) in the presence of ^32^P-dATP for 1h at RT. Analysis was performed by spot assay as described in Material and Methods. The y-axis represents the percentage of radioactive DNA synthesis expressed in % of full plasmid replication. Addition of Aphidicolin to NPE (150 uM final) more severely reduces incorporation in MNU+, MNU++ and ENU plasmids, compared to control and MMS plasmids. Fold reduction in incorporation under aphidicolin conditions is shown.

**Fig. S3.**
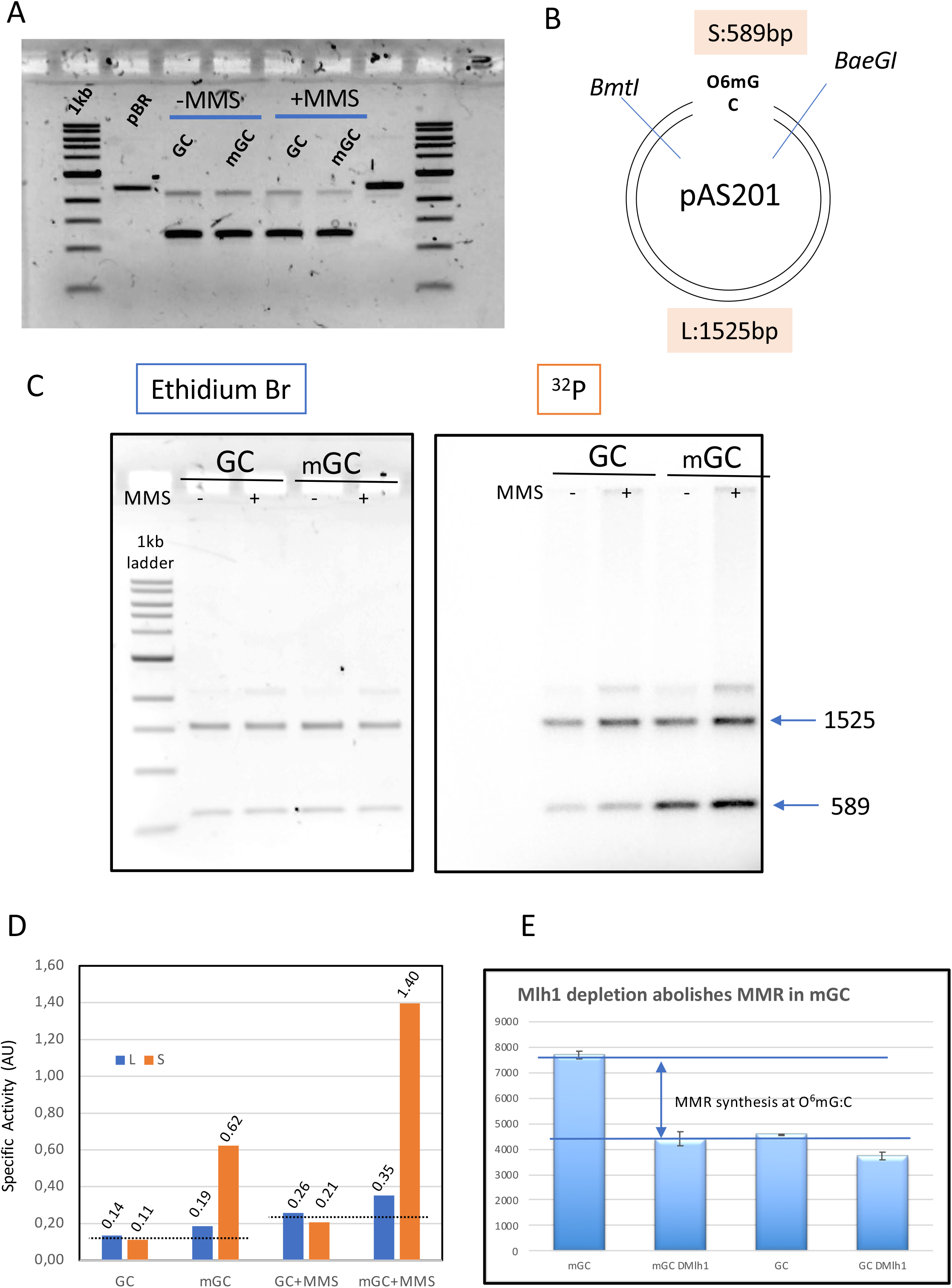
Mapping repair synthesis in the vicinity of a single O^6^mG adduct. **A:** Agarose gel electrophoresis of the native form of GC, mGC, GC+MMS and mGC+MMS **B**. Restriction map of pAS201, the two single cutter enzymes *BmtI* and *BaeGI* yield a short S (589bp) and long L (1525bp) fragment, respectively. Fragment S contains the single O^6^mG adduct. **C**. Plasmids GC, mGC, GC+MMS and mGC+MMS were incubated in NPE in the presence of α^32^P-dATP and extracted after 2h. The purified plasmids were digested with restriction enzymes *BmtI* and *BaeGI*. Digested plasmids were analyzed by agarose gel electrophoresis and visualized by ethidium bromide staining and by ^32^P-imaging. **D**. The specific activity (SA) of a given fragment (S or L) was quantified as the amount of ^32^P incorporated divided by the amount of DNA deduced from the Ethidium bromide fluorescence image (expressed in arbitrary units AU). In control plasmid GC, a similar SA value was observed for both fragments as expected for residual background incorporation; similarly, in MMS treated plasmid (GC + MMS), the SA value is similar for S and L fragments. The slightly increased SA value in GC+MMS compared to GC is compatible with repair synthesis by BER at randomly located MMS-induced lesions. For plasmids containing a single O^6^mG:C lesion (mGC and mGC+MMS), there is a robust increase in SA of the short compared to the long fragments. Incorporation, above background, due to O^6^mG, in the absence of MMS, amounts to 0.065 and 0.495 AU for long and short fragments, respectively (signal above dotted line in S3D). Similarly, in the context of MMS lesions, incorporation, above background, due to O^6^mG amounts to 0.115 and 1,17 AU for long and short fragments, respectively (signal above dotted line in S3D). Taken together these results clearly show that O^6^mGC-mediated repair specifically takes place in the S fragment, with only modest spill-over into the L fragment (10-15%). **E**. Plasmid with a single O^6^mG:C lesion site (mGC) and control plasmid (GC) were incubated in NPE depleted with anti Mlh1 antibodies and mock treated NPE. Incorporation was measured into whole plasmids. The increase in incorporation into mGC compared to GC observed upon incubation in NPE fully disappeared when Mlh1 was depleted. This experiment shows that the increase in incorporation observed in mGC compared to incorporation in control GC plasmid can specifically be attributed to mismatch repair activity at the single O^6^mG:C lesion.

**Fig. S4.**
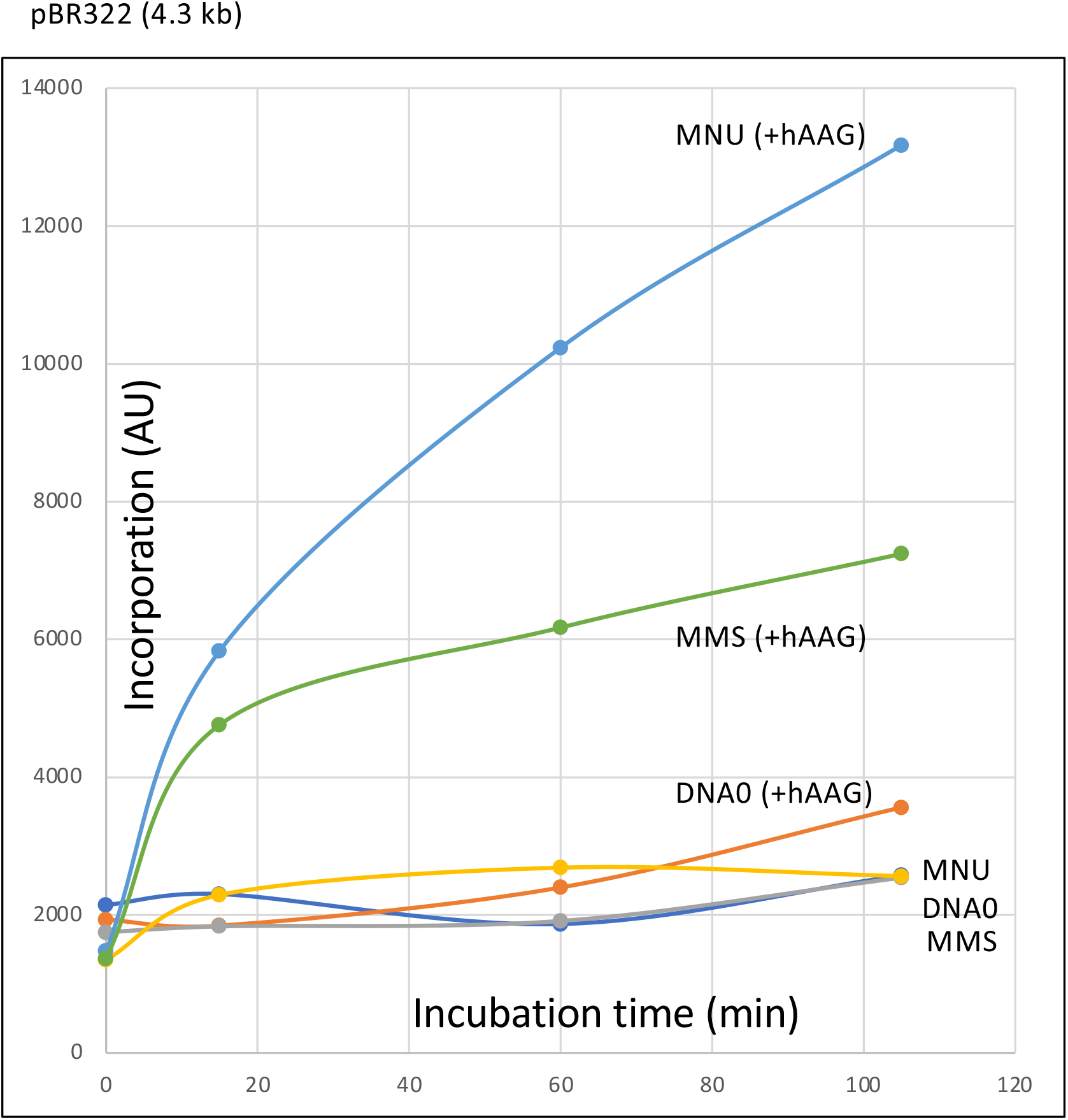
DNA repair synthesis in HSS extracts depends upon addition of AAG glycosylase (NEB, Biolabs). Plasmid pBR322, unmodified (DNA0) or modified with -MMS or -MNU to a similar extent (≈1 N-alkyl adduct/500nt), were incubated in the presence of α^32^P-dATP. Repair synthesis was monitored at room temperature as a function of time using the spot assay described above (Fig. 1A). Lack of repair synthesis in HSS extracts, is restored upon addition of hAAG glycosylase (150nM).

**Fig. S5.**
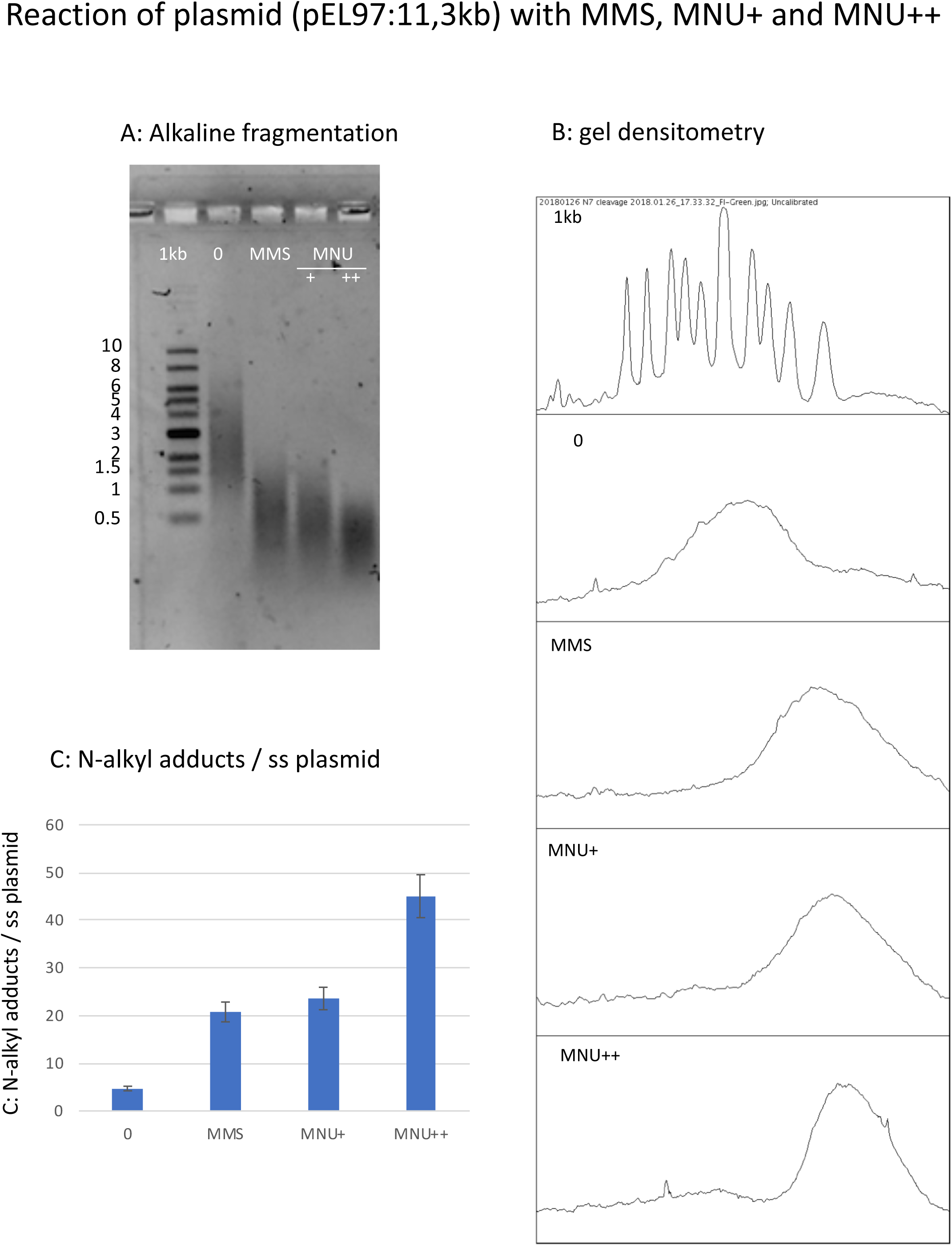
Estimation of N-alkylation levels of modification by MMS and MNU. A. Alkaline fragmentation of alkylated plasmid (pEL97, 11.3 kb) DNA as outlined in Fig. S1A analyzed by agarose gel electrophoresis. B. Densitometry of the agarose gel. C. Estimation of the average number of alkali cleavage sites per plasmid strand. The data reveal that the average number of cleavage sites per plasmid strand is 21-23 for both MMS and MNU+ reaction conditions (⇔ the average distance between two sites is ≈ 510 nt). For MNU++, the average distance between two cleavage sites is ≈254 nt.

**Fig. S6.**
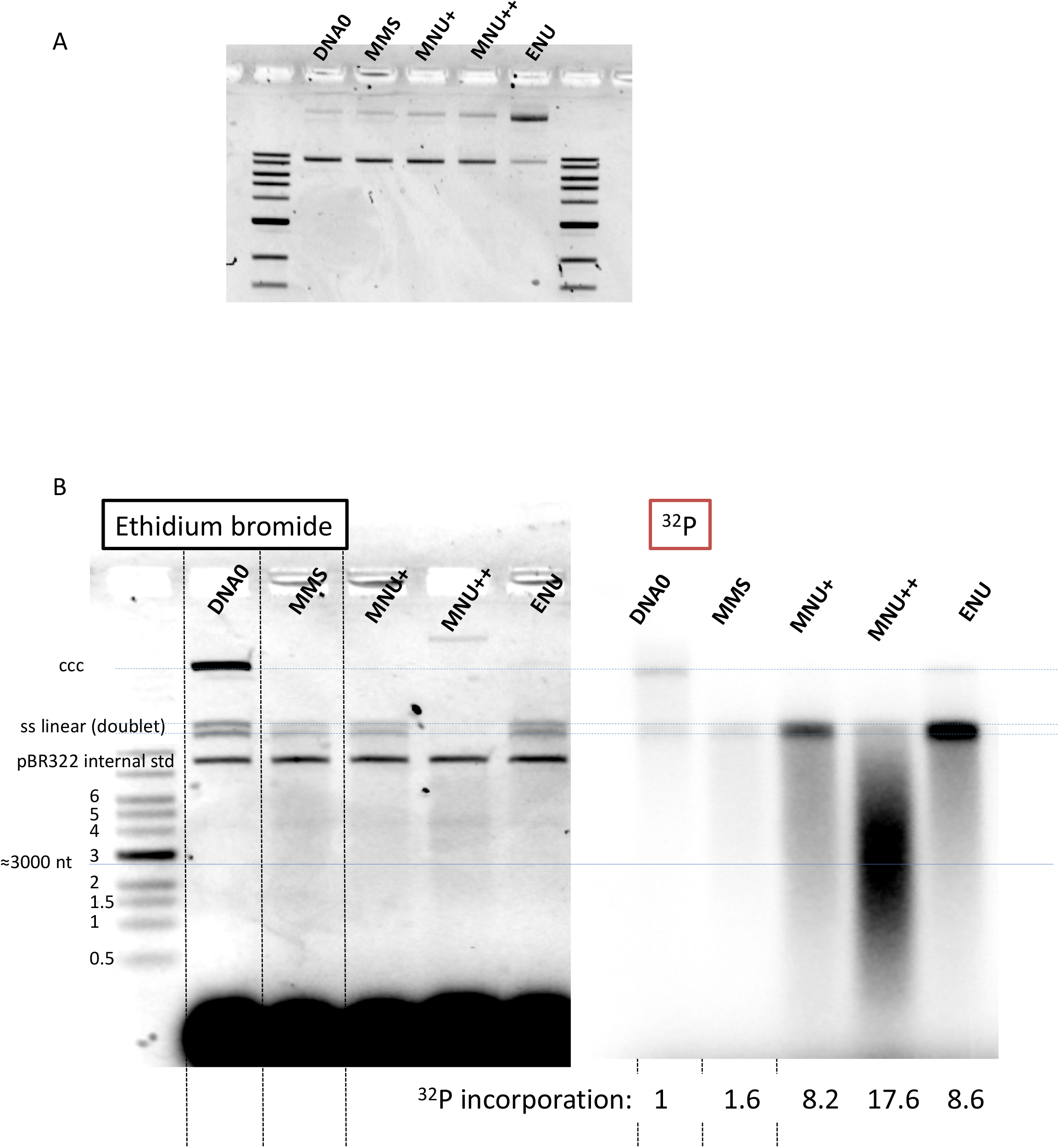
Fragmentation of alkylated plasmid as analyzed on AGE loaded under alkaline conditions. **A:** Alkylation did not affect plasmids topology except for ENU treatment that increases relaxation as seen in the AGE image. **B:** Analysis by AGE, under alkaline loading conditions, of alkylated plasmids (pEL97: 11,3kb) incubated in NPE in the presence of α^32^P-dATP (same samples as in Fig. 4A). Loading under alkaline conditions allowed single-stranded nicks to be revealed. In the different lanes, an internal standard band, pBR (ccc), was introduced before the extraction procedure in order to assess consistent recovery of DNA during extraction (as seen by ethidium bromide staining). For the alkylated plasmids, incubation in NPE led to massive conversion of plasmid DNA into a smear with a low amount of DNA present as a single-stranded linear band (doublet). For the MNU++ sample, no linear ss DNA was seen, all DNA was converted into a smear. The smearing can best be seen in the ^32^P image. We suggest that the ss DNA fragments that form the smear arise between a gap caused by a MMR event at a O^6^mG sites and an uncompleted BER event at a neighbor lesion. It cannot be excluded that some fragments result from MMR attempts at two O^6^mG lesions. The intense ^32^P labelling of the fragments in the smear result from incorporation of up to several hundred nt during a single MMR repair synthesis. Relative ^32^P incorporation normalized to 1 for DNA0 are indicated below the ^32^P-image.

